# Identification of structural features of selected flavonoids responsible for neuroprotection using a *Drosophila* model of Parkinson’s disease

**DOI:** 10.1101/2022.06.03.494711

**Authors:** Urmila Maitra, John Conger, Mary Magdalene (Maggie) Owens, Lukasz Ciesla

**Author notes:** Co-corresponding authors: Dr. Urmila Maitra and Dr. Lukasz Ciesla, Department of Biological Sciences, University of Alabama, 2320 Science and Engineering Complex, Tuscaloosa, Alabama 35487-0344.

## Abstract

Nature-derived bioactive compounds have emerged as promising candidates for the prevention and treatment of diverse chronic illnesses, including neurodegenerative diseases. However, the exact molecular mechanisms underlying their neuroprotective effects remain unclear. Most studies focus solely on the antioxidant activities of natural products which translate to poor outcome in clinical trials. Current therapies against neurodegeneration only provide symptomatic relief thereby underscoring the need for novel strategies to combat disease onset and progression. We have employed an environmental toxin-induced *Drosophila* Parkinson’s disease (PD) model as an inexpensive *in vivo* screening platform to explore neuroprotective potential of selected dietary flavonoids. We have identified a specific group of flavonoids known as flavones displaying protection against paraquat (PQ)-induced neurodegenerative phenotypes, involving reduced survival, mobility defects and enhanced oxidative stress. Interestingly, the other groups of investigated flavonoids, namely, the flavonones and flavonols failed to provide protection indicating a requirement of specific structural features that confer protection against PQ-mediated neurotoxicity in *Drosophila*. Based on our screen, the neuroprotective flavones lack a functional group substitution at the C3 and contain α,β-unsaturated carbonyl group. Furthermore, flavones-mediated neuroprotection is not solely dependent on antioxidant properties but also involves regulation of neuroinflammatory responses. Our data identify specific structural features of selected flavonoids that provide neuroprotection against environmental toxin-induced PD pathogenesis that can be explored for novel therapeutic interventions.

**Graphical Abstract:** 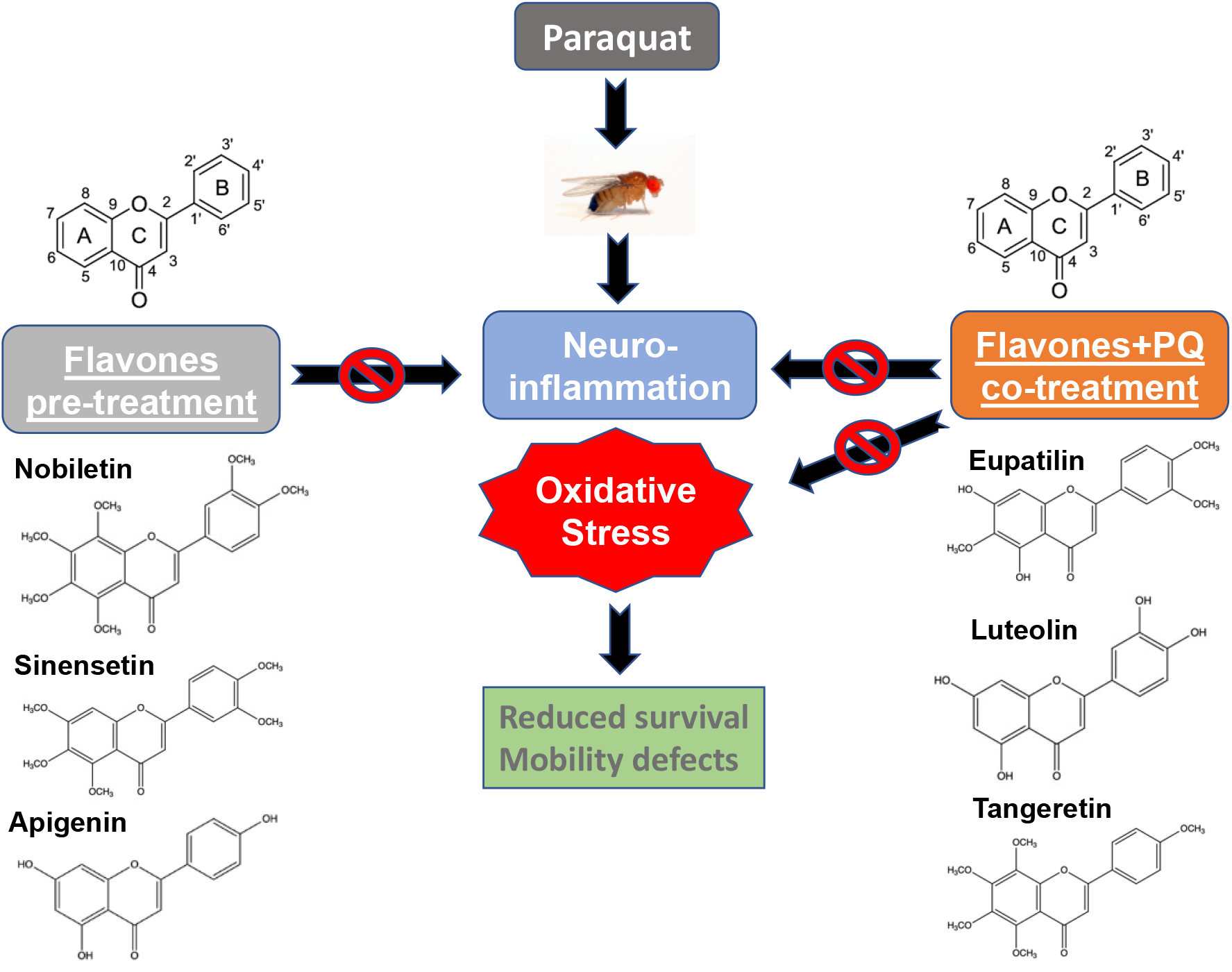

## Introduction

The incidence of chronic metabolic non-communicable ailments such as neurodegenerative diseases (ND) has been on the rise and become one of the most challenging global threats and economic burdens of the 21^st^ century (1). Currently, there are no treatment options available to cure NDs, and the approved therapies only manage disease symptoms (2). A score of epidemiological studies strongly points to the fact that lifestyle modifications such as regular physical activity and/or following certain types of diet, such as the Mediterranean diet, have been linked to a lower risk of developing Alzheimer’s (AD) or Parkinson’s disease (PD) (3-11). Bioactive compounds, often referred to as specialized (or secondary) metabolites abundant in plant-rich diets, have been linked to beneficial and disease-preventive effects of dietary approaches (12). Polyphenols belong to a class of abundant specialized metabolites that have been often considered as majorly responsible for the beneficial effects of plant-based diets (2, 12, 13). Numerous recent epidemiological studies have shown that people consuming a diet rich in flavonoids, a subclass of polyphenolic compounds have been less likely to develop AD or PD (14-16).

Flavonoids are commonly present in plants and have received massive attention in the scientific literature, including the investigation of neuroprotective properties (1, 2). However, the exact molecular mechanism of action of flavonoids providing neuroprotection remains elusive (1, 12). For the past fifty years, the predominant hypothesis in the field promulgated the antioxidant activities of polyphenols as majorly responsible for their beneficial effects (1). However numerous *in vivo* studies, including clinical trials disproved direct antioxidant activity of polyphenols (1). Additionally, the upregulation of the antioxidant response element signaling pathway involving the Nuclear factor erythroid 2-related factor 2 (Nrf2) has also been shown as insufficient to provide neuroprotection in certain animal models of NDs (17).

Some have proposed that flavonoids may have evolved in plants as deterrents to provide protection from herbivores and omnivores (1). Mattson et al. suggested that flavonoids, and other specialized metabolites at low concentrations activate numerous cellular stress response pathways, proposing the hormetic mechanism of action (1, 18-22). However, many polyphenols remain safe at high concentrations failing to follow the hormesis principle. Sinclair et al. in the xenohormesis hypothesis suggested that heterotrophs sense the increasing levels of specialized metabolites in stressed autotrophs that allow them to prepare in advance for different types of environmental adversities (23-32). Both hypotheses also assume all phytochemicals follow the hormetic patterns disregarding the evident differences between various classes of specialized metabolites, such as alkaloids, terpenoids or polyphenols.

Despite many years of research on flavonoids, none of the investigated compounds has been successfully registered as a drug (2, 12). This has led to the classification of these compounds as invalid/improbable metabolic panaceas (IMPs) lacking drug-like characteristics, such as an exquisitely specific interaction with a definable receptor (2, 12, 33). Additionally, flavonoids have been often classified as pan-assay interference compounds (PAINs) due to their ability to interact with numerous targets in *in vitro* assays (34). Many essential nutrients, such as vitamins or essential fatty acids (formerly known as vitamin F) play important roles in biological systems and fail to meet the reductionist definition of a drug (2, 12). NDs are considered multifactorial diseases caused by complex interactions between genetic and environmental factors (35, 36). Single compounds displaying pleiotropic effects, such as flavonoids, have been suggested to be more effective in the prevention or treatment of multifactorial diseases (2). Seigler et al. proposed that flavonoids and coumarins may play an essential role in human nutrition, following the concept of vitamin P, proposed by Szent-Györgyi (12).

Previous structure-activity relationship studies focused majorly on flavonoid antioxidant effects tested using *in vitro* assays (37, 38). There is only very limited number of published studies focusing on understanding how structure of flavonoids impacts their biological effects in the context of a living organism. Many biologically active flavonoids contain electrophilic α,β-unsaturated carbonyl group similarly to the oxidation products of essential omega-3 fatty acids (2). The presence of highly reactive electrophilic group in flavonoids used at supraphysiological concentrations in many *in vitro* assays, may be responsible for the false positive results reported in the literature. Therefore, it is of crucial importance to investigate the biological activity of flavonoids, including interactions with multiple targets, in the context of a living organism at physiologically relevant concentrations.

We have employed a well-established paraquat-induced *Drosophila* model to screen selected flavonoids against PD symptoms in the context of a living organism (39). This fly model mimics the clinical hallmarks of PD in humans, including motor deficits, increased neuroinflammation and oxidative stress and serves as an inexpensive platform to screen plant derived flavonoids through diet (2, 40). The flavonoids were selected based on their structural similarity to gardenin A, which we previously identified as neuroprotective in this *Drosophila* model of PD (17). We investigated the neuroprotective potential of a group of selected flavonoids in our attempt to identify the structural elements linked to neuroprotection and decipher the possible molecular mechanism of action of these compounds.

## Materials and methods

### Drosophila culture and stocks

The wild-type strain *Canton S* was purchased from the Bloomington Drosophila Stock Center at Indiana University and used in all experiments. The fly stocks were raised on standard Nutri-fly BF medium (Genesee Scientific) containing cornmeal, corn syrup, yeast, and agar at 25 °C under a 12 h of light and 12 h of darkness cycle. Age-matched male flies three to five days old post-eclosion were used for all assays.

### Chemicals

The flavonoids used in this study and the following chemicals were purchased from Sigma-Aldrich: Tangeretin Catalog# T8951 (>95% purity), Nobiletin N1538 (>97% purity), Apigenin 42251 (>95%), Sinensetin SML1787 (>98%), Luteolin 72511, Fisetin PHL82542, Eupatilin SML1689 (>98%), Naringenin 52186 (>95%), Hesperetin PHL89222, Kaempferol 96353, Quercetin Q4951 (>95%), Myricetin 70050 (>96%), Naringin 71162 (95%), Hesperidin H5254, Sucrose, and paraquat (methyl viologen dichloride hydrate). TRIzol reagent and the primers were obtained from Thermo Fisher Scientific. The Direct-zol RNA microprep kits for total RNA isolation were purchased from Zymo Research and the iQ SYBR Green Supermix was from Biorad.

### Drosophila feeding regimen

Adult male flies aged 3–5 days post-eclosion were maintained in vials (10/vial) and fed daily on filter paper saturated with specified concentrations of paraquat with or without flavonoids in 2.5% sucrose or with 2.5% sucrose only. Mortality was monitored daily until all flies were dead. Survival assays were performed with at least ten independent biological replicates of 10 males each for different feeding conditions.

*CantonS* flies were fed either 2.5% sucrose or specified concentrations of different flavonoids containing the blue food dye (1% FD&C Blue#1) to measure food intake quantitatively. The flies were washed and homogenized in 1X PBS containing 1% Triton X-100 followed by centrifugation and the supernatants were measured at OD 630 nm. Data were analyzed using the two-tailed *Student’s* t-test and error bars indicate the standard deviation.

### Mobility assay

Negative geotaxis assays were performed to detect the mobility defects in adult male flies from different feeding group. Ten flies per feeding condition were placed in an empty plastic vial and gently tapped to the bottom. The percentage of flies that crossed a line 5 cm from the bottom of the vial in 20 sec were recorded to compare between different feeding conditions. Five independent biological replicates were assayed three times at 5 mins intervals and the average percentages were calculated and plotted. Statistical significance between different feeding conditions was calculated using one-way analysis of variance (ANOVA) for *P* < 0.05.

### Quantitative real time RT-PCR

Total RNA was extracted from the heads of 25-30 adult male flies using TRIzol Reagent and Direct-zol RNA microprep kit (Zymo Research) followed by cDNA preparation from 0.5–1 μg total RNA using the High Capacity cDNA Reverse Transcription kit (Applied Biosystems). The resulting cDNA samples were diluted, and specific transcripts were detected using the iQ SYBR Green Supermix (Biorad) in a StepOnePlus Real time PCR System according to the manufacturer’s protocols. The relative levels of specified transcripts were calculated using the ΔΔCt method, and the results were normalized based on the expression of *ribosomal protein L32* (*RpL32/RP49*) as the endogenous control within the same experimental setting. At least three independent biological replicates were performed in response to specified feeding conditions and compared to the control sucrose-fed flies (assigned a value of 1) for data analysis.

### Statistical analysis

Data were analyzed using the GraphPad Prism 8 software (GraphPad Software, Inc., La Jolla, CA). Statistical significances of gene expression between different groups were determined using a non-parametric Mann Whitney U-test. For survival and mobility assays, log-rank test and one-way analysis of variance (ANOVA) were used respectively, to compare the differences between specified feeding groups. Results are expressed as mean ± SEM, and *p* < 0.05 was considered statistically significant. Details of the specific analysis are mentioned in the figure legends.

## Results

### A specific group of flavonoids known as flavones exert protection against paraquat-induced toxicity in a *Drosophila* PD model

We employed an environmental toxin-induced *Drosophila* PD model as a screening platform to identify flavonoids with neuroprotective potential. Exposure to the herbicide paraquat (PQ) has been previously shown to exhibit reduced survival, impaired climbing abilities, and dopaminergic neuron loss in the wild-type *Drosophila* strain, *Canton S* (39, 41-43). Adult male flies were used in this study due to their higher sensitivity to PQ toxicity leading to PD phenotypes earlier than female flies, thereby mimicking the higher prevalence of PD in human male patients (41). Adult male *Canton S* wild type flies were fed 5 mM PQ dissolved in 2.5 % sucrose solution on filter paper to induce parkinsonian symptoms, including reduced survival and enhanced mobility defects. This *Drosophila* PD model was used to screen three groups of flavonoids, namely flavones, flavonones and flavonols as listed in Table 1. Flavonoids consist of different groups of polyphenolic compounds that are ubiquitously present in plants and one of the most common groups of phytochemicals found in the human diet (44). Flavonoids are subdivided into different subgroups depending on the presence of methoxy or hydroxy groups, the degree of unsaturation and oxidation status of the C-ring (44, 45). Flavones have a double bond between positions 2 and 3 and a ketone in the position 4 of the C-ring. Some flavones tested in this model are polymethoxylated. Flavonols have a hydroxyl group in the position 3 of the C-ring and they are diverse in methylation and hydroxylation patterns. On the other hand, flavonones have the C-ring saturated between positions 2 and 3 as compared to flavones (45, 46). The flavonoids were selected that are structurally related to each other but differ in the functional groups attached to the parent flavonoid skeleton and unsaturation of the C-ring to elucidate the structural features that confer protection against PQ-induced toxicity using survival assays.

**Table 1.**
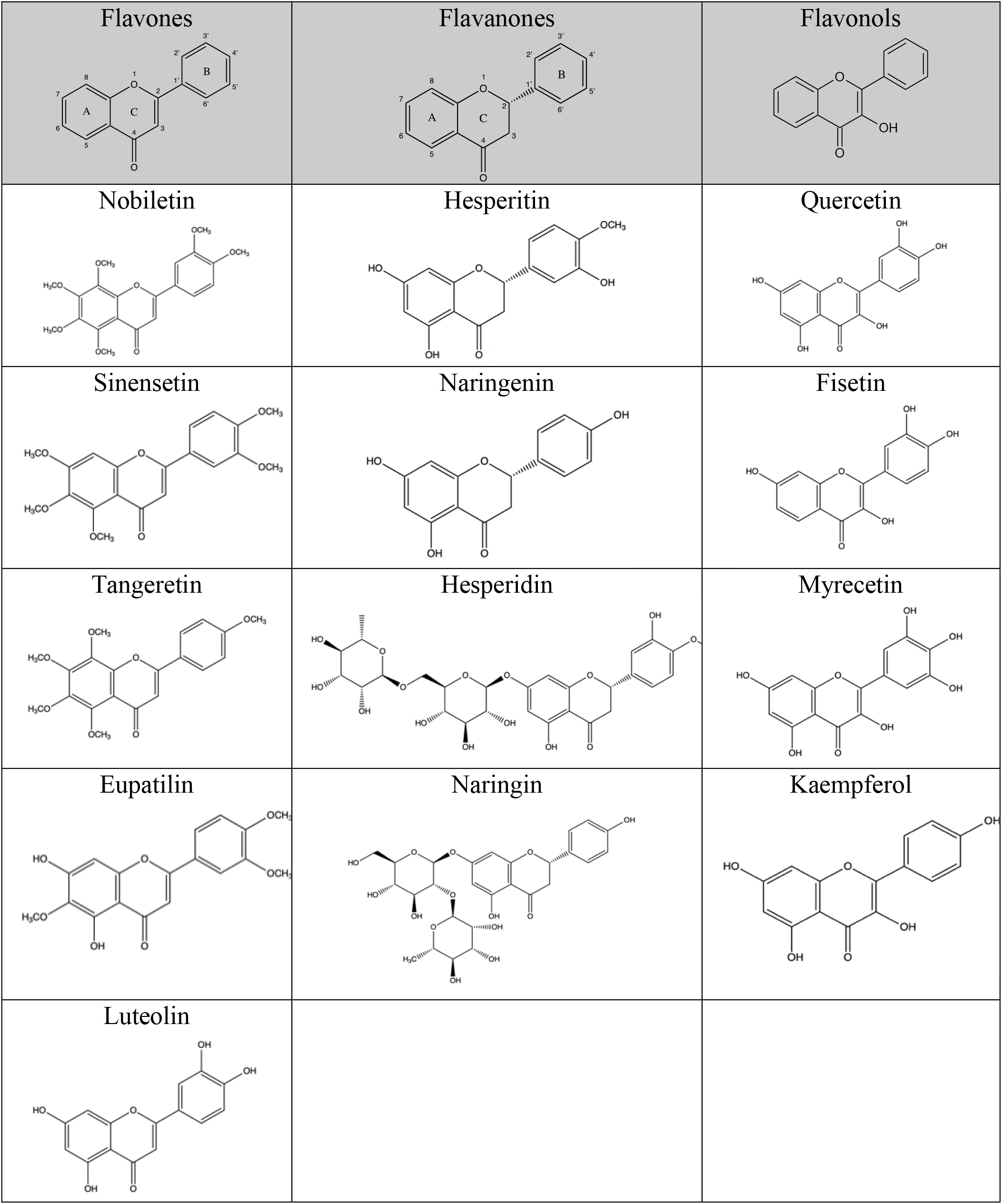
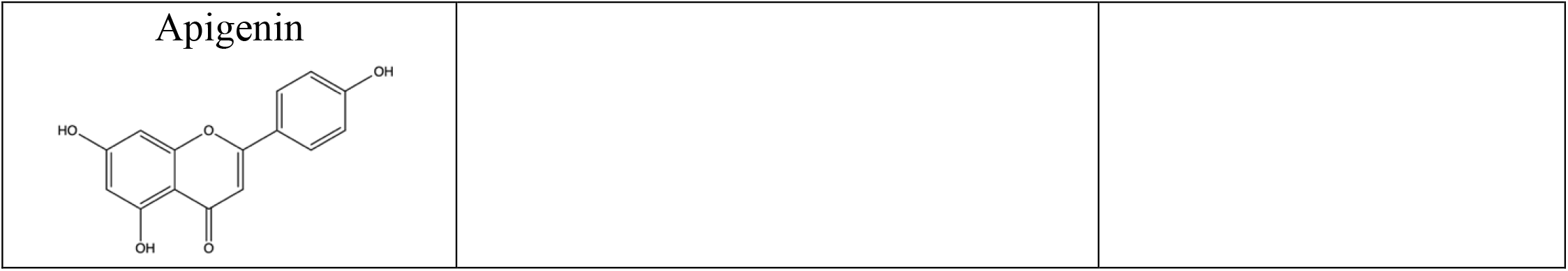
Structures of the investigated flavonoids.

We used two modes of feeding regime, Group I: pre-treatment with flavonoids for 4 days before continuous exposure to PQ and Group II: co-treatment with flavonoids along with PQ as shown in Fig.1A. For Group I, adult male flies were pre-fed for 4 days with either 2.5% sucrose solution as control or specified flavones alone diluted in 2.5% sucrose solution, and then exposed to 5 mM PQ continuously and survival was scored every 24 h post-exposure, and for Group II, the flies were co-fed with flavonoids diluted in 2.5% sucrose solution along with 5 mM PQ. The duration of 4 days of pre-treatment was chosen based on our earlier findings that showed neuroprotection by another polymethoxy flavonoid, Gardenin A against PQ toxicity using the *Drosophila* PD model (17). We selected the specified concentrations of the flavonoids ranging from 5-100 µM after stringent preliminary screening that gave optimal reproducible data in survival assays and were considered physiologically relevant in the micromolar concentration ranges.

**Figure 1.**
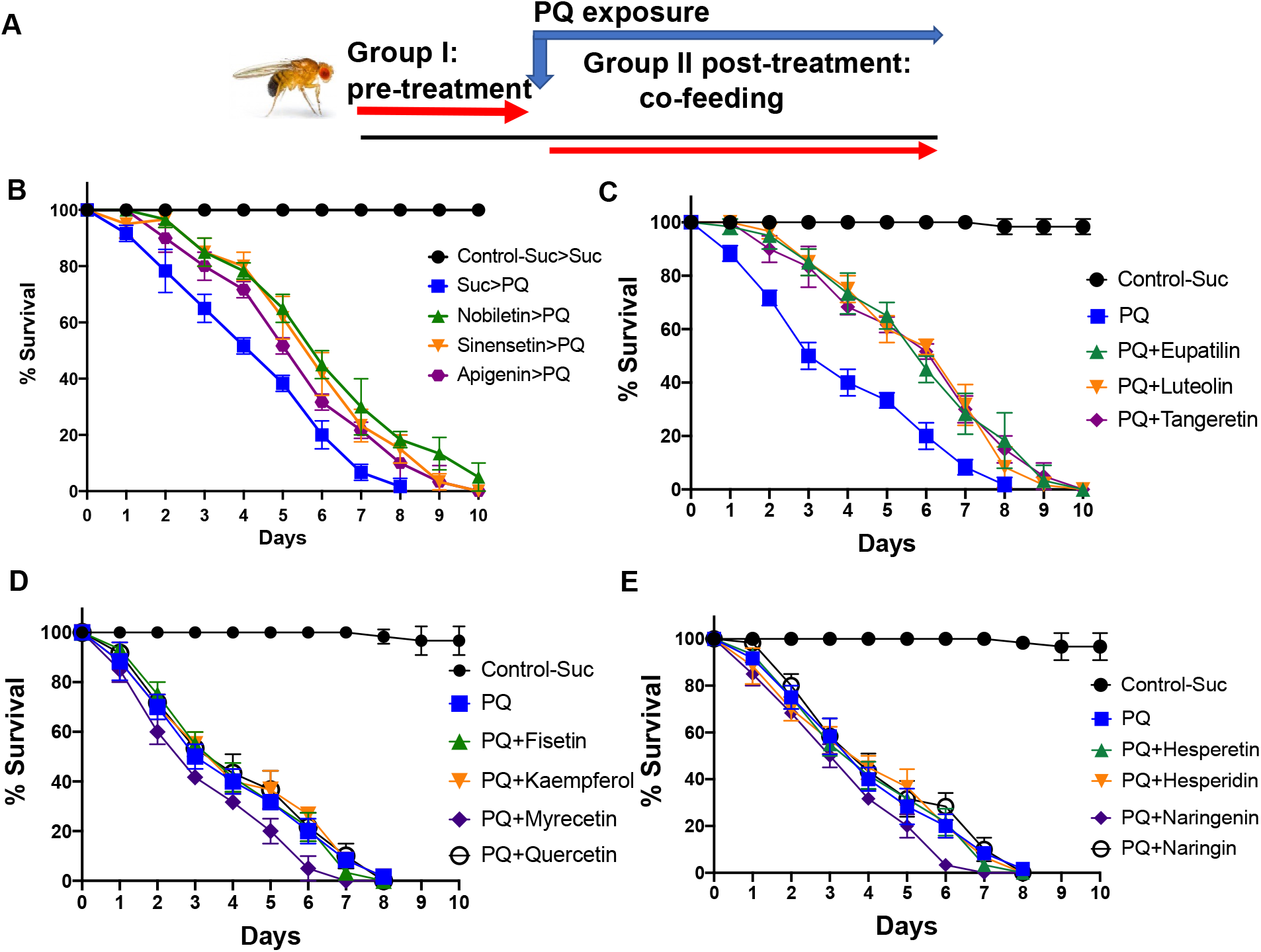
Effects of different groups of flavonoids against paraquat-induced toxicity in a *Drosophila* PD model. A. Schematic representation of the pre- and co-feeding regimen. In the pre-treatment mode, adult male flies were either pre-fed for 4 days with 2.5% sucrose or specified flavonoids followed by continuous exposure to 5 mM PQ and were scored daily for survival. In the co-treatment mode, adult male flies were exposed to both flavonoids and 5 mM PQ at the same time and were scored daily for survival. Survival data are representative of ten independent biological replicates with 10 male flies per feeding conditions. B. Survival assays were performed using adult male flies pre-fed with sucrose or specified flavones (10 µM nobiletin, 5 µM sinensetin and 10 µM apigenin) followed by continuous exposure to 5 mM PQ as outlined above and the number of live flies was recorded every 24 h until all of the flies were dead and the average survival percentages were plotted. Statistical significances between the PQ-fed and the flavone pre-fed groups followed by PQ exposure were determined using the log-rank test (Suc>PQ vs Nobiletin>PQ, Suc>PQ vs Sinensetin>PQ ***p<0.001; Suc>PQ vs Apigenin>PQ **p<0.01). C. Survival assays were performed using the co-feeding mode in which adult male flies were exposed to both 5 mM PQ or specified flavones (5 µM eupatilin, 100 µM luteolin and 20 µM tangeretin) and the number of live flies was recorded every 24 h and the average survival percentages were plotted. Statistical significances between the PQ-fed and the co-fed PQ+flavone groups were determined using the log-rank test ***p<0.001. D. Survival assays were performed using the co-feeding mode in which adult male flies were exposed to both 5 mM PQ or specified flavonols (25 µM fisetin, 20 µM kaempferol, 25 µM myricetin, 25 µM quercetin) and the number of live flies was recorded every 24 h and the average survival percentages were plotted. No statistical significances between the PQ-fed and the flavonol-fed groups were observed using the log-rank test. E. Survival assays were performed using the co-feeding mode in which adult male flies were exposed to both 5 mM PQ or specified flavonones (20 µM hesperitin, 20 µM hesperidin, 20 µM naringenin, 25 µM naringin) and the average survival percentages were plotted. No statistical significances between the PQ-fed and the flavonone-fed groups were observed using the log-rank test.

Consistent with earlier findings, exposure to PQ leads to a gradual increase in mortality (Fig.1B) (39). However, the group of flavones, nobiletin, sinensetin and apigenin conferred protection when pre-fed for 4 days prior to PQ exposure. Interestingly, nobiletin and sinensetin were protective when the flies were pre-exposed to flavones intermittently for 4 days (flavones on Days 1 and 3; sucrose on Days 2 and 4 for the pre-treatment duration) while apigenin improved survival against PQ-toxicity when pre-fed continuously for 4 days prior to PQ exposure as depicted in Fig. 1B. In contrast, the other group of flavones, eupatilin, luteolin and tangeretin were protective against PQ-induced toxicity only in co-feeding mode using survival assays as shown in Fig.1C. In both pre-and co-treatment groups, the flavones significantly improved the median survival to PQ and extended the survival rate by 1-3 days as compared to only PQ-treated groups (Figs. 1B&C).

The same feeding regimen were followed for the group of flavonones (hesperitin, hesperidin, naringenin, naringin) and flavonols (fisetin, kaempferol, myricetin, quercetin) to identify compounds that were protective against PQ-induced toxicity in either pre-feeding or co-feeding modes of treatment. Flavonoids can be found in plants in both glycoside-bound and free aglycon forms. Both flavonone glycosides (hesperidin and naringin) and aglycones (hesperetin and naringenin) were included in the screen to evaluate the effects on PQ-mediated morbidity in flies. As shown in Figs.1D and E, the data suggest that none of the selected compounds in the groups of flavonones and flavonols were able to rescue against PQ-induced toxicity in survival assays using both pre- and co-feeding modes of treatment. Additionally, we observed that only aglycones exert protective effects in our model.

Feeding assays rely on the amount of food consumed by the flies and the taste and odor of different flavonoids mixed with the control food might influence the food intake ability. Therefore, we performed feeding assays using the dye (1% FD&C Blue#1) mixed with food to ensure that exposure to different flavonoids do not lead to significant differences in the food intake abilities as compared to the sucrose-fed controls. Our data confirm comparable levels of food consumption in flies exposed to different groups of flavonoids as shown in Fig.2 A and B. Overall, the results suggest that the specific group of flavonoids known as flavones confer protection against PQ toxicity in *Drosophila*.

**Figure 2.**
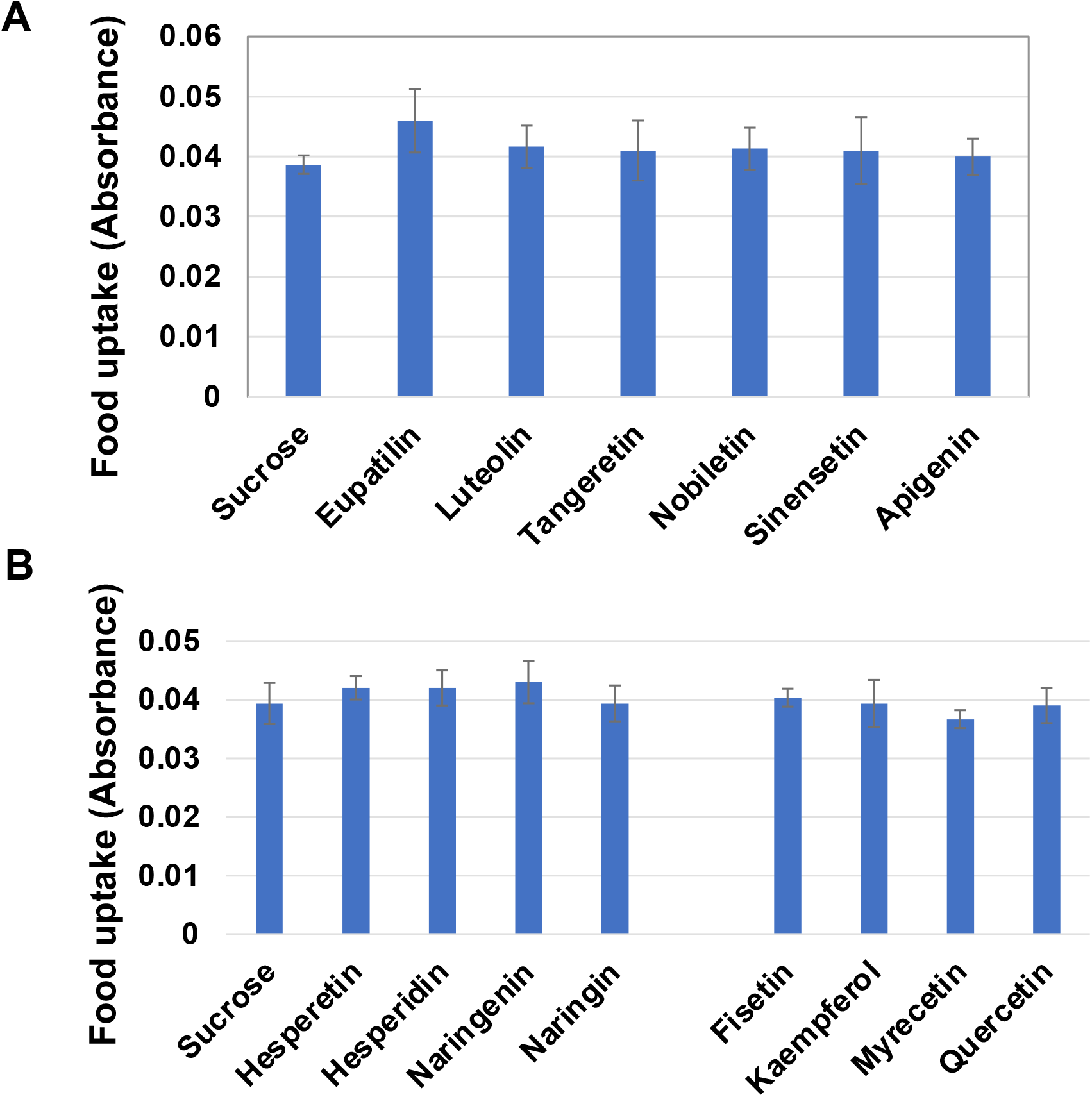
Feeding assays to quantitate food uptake upon exposure to different flavonoids. Adult male flies were fed either sucrose or specified flavonoids A. flavones B. flavonones or flavonols mixed with the blue food dye (1% FD&C Blue#1) and the dye content was measured spectrophotometrically at 630 nm. Error bars indicate the standard deviation. No statistical significances between the control sucrose-fed group and the specified flavonoid-fed groups were observed using the Student’s t-test. p>0.05.

### Identification of specific structural features of flavonoids required to rescue PQ-induced mobility defects

Our next goal was to determine the effect of different groups of flavonoids on PQ-mediated locomotion defects using negative geotaxis assay. Severe locomotion impairment is a common symptom of PD caused by the deterioration of dopaminergic neurons in the midbrain (39, 41). We used the same feeding regime as specified in Fig.1A and climbing assessment was performed at 48 h time point post PQ exposure. The flies were either pre-fed for 4 days with sucrose or different concentrations of flavonoids as specified followed by continuous exposure to PQ or co-fed with PQ and climbing assessment was performed at 48 h time points post-exposure. As shown in Fig. 3A, PQ exposure significantly reduced climbing abilities to 52% as compared to the 92% of the sucrose-fed control group. However, pre-treatment with specific group of flavones, including, nobiletin, sinensetin and apigenin, significantly restored mobility impairment after PQ exposure. Similarly, co-feeding PQ with the flavones, eupatilin, luteolin and tangeretin also significantly improved PQ-induced mobility defects as shown in Fig. 3B. However, the structurally related flavonoids, flavonones (hesperitin, hesperidin, naringenin, naringin) and flavonols (fisetin, kaempferol, myricetin, quercetin) failed to improve mobility defects in flies exposed to PQ (Fig. 3C and D). Our data suggest that certain structural features of flavones including the presence of a double bond at the C2-C3 position (α,β-unsaturated carbonyl) of the flavonoid skeleton and the lack of a functional group substitution at the C3 position are essential to confer protection against PQ-mediated toxicity in *Drosophila*. The compounds belonging to the flavonones and flavonols subgroups do not comply with these structural features and thus fail to provide protection against PQ exposure. Our findings confirm that only specific group of flavonoids known as flavones are protective against paraquat-induced toxicity and mobility defects, which could potentially be attributed to the structural features like the degree of unsaturation and functional groups attached to the C-ring of the parent flavonoid structure.

**Figure 3.**
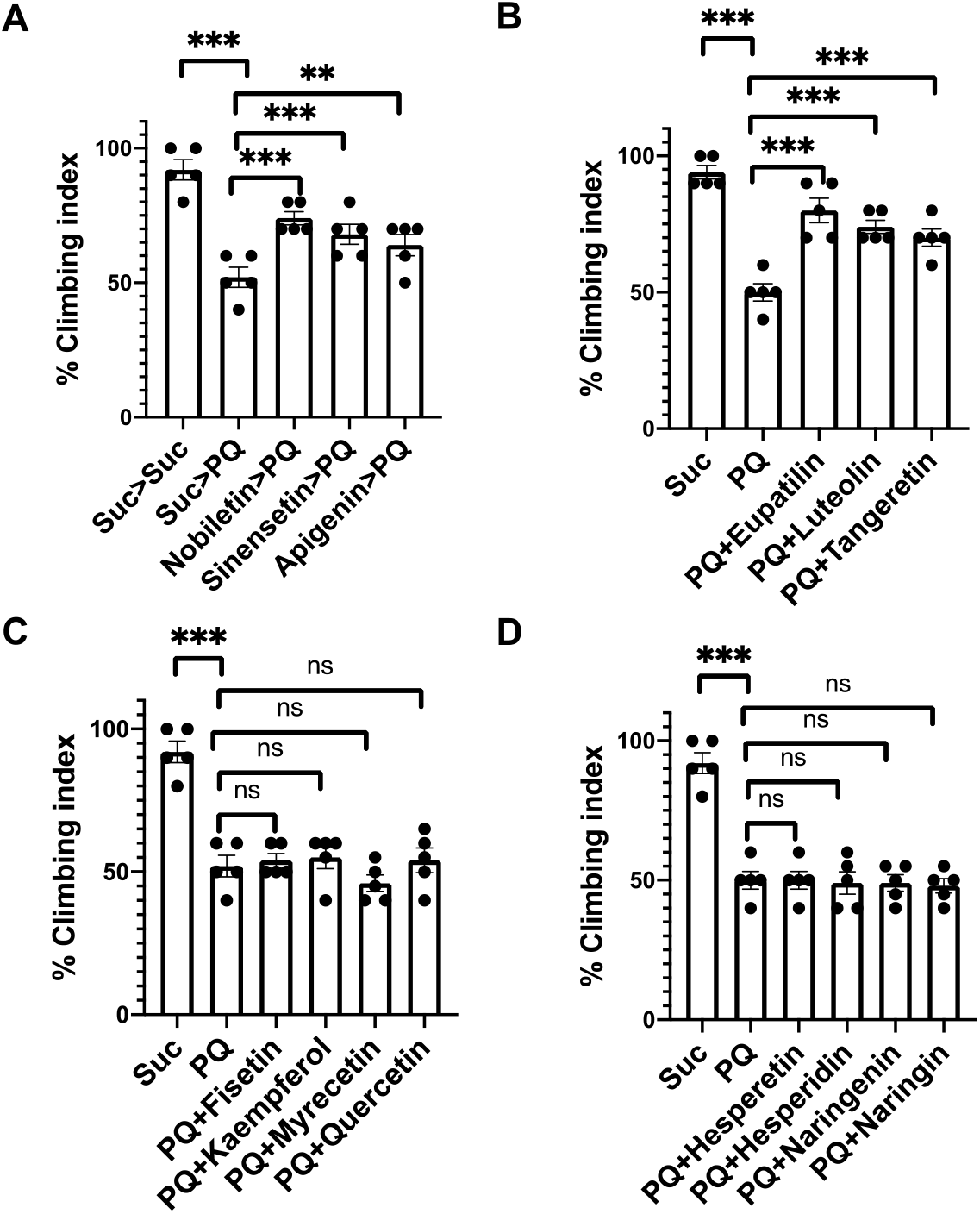
Effects of different groups of flavonoids against paraquat-induced mobility defects. Negative geotaxis assays were used to determine the effect of flavonoids on the climbing abilities of flies exposed to PQ. The number of flies able to cross 5 cm within 20 s were recorded and plotted at 48 h. Data are representative of at least five independent experiments with 10 male flies per group. A. Adult male flies were either pre-fed for 4 days with 2.5% sucrose or flavones (nobiletin, sinensetin and apigenin) followed by continuous exposure to 5 mM PQ and the % climbing index were plotted at 48 h. B. Mobility assays were performed using the co-feeding mode in which adult male flies were exposed to both 5 mM PQ or specified flavones (eupatilin, luteolin and tangeretin) and data were analyzed at 48 h. The protective effects of specified flavonoids on the climbing abilities of flies exposed to PQ were determined by mobility assays. C. flavonols (fisetin, kaempferol, myricetin, quercetin) and D. flavonones (hesperitin, hesperidin, naringenin, naringin). **p < 0.01; ***p < 0.001 based on one-way ANOVA between indicated feeding conditions.

### The antioxidant properties of flavones are not sufficient to confer protection against PQ-induced oxidative stress

Increased oxidative stress has been linked to the onset and progression of neurodegenerative diseases (ND), including AD and PD (17, 47). Exposure to the herbicide PQ is known to induce oxidative stress leading to the clinical symptoms of PD in both mammalian and invertebrate models (17, 48). Flavonoids have been shown to reduce oxidative stress by inducing the antioxidant response through the nuclear factor erythroid 2–related factor 2 (Nrf2), which regulates the expression of several downstream antioxidant genes and detoxifying proteins such as thioredoxins, glutathione synthetase, and glutathione S-transferases to counterbalance the accumulation of free radicals within the cells (46, 49). Nrf2 homologs belong to the *cap‘n’collar (cnc)* subfamily of leucine zippers and known as the *cnc* gene in *Drosophila* (50). Therefore, we evaluated the effect of flavones that were protective in either pre- or co-feeding modes on the genes involved in the antioxidant response pathways. The flies were pre-fed for 4 days with either sucrose or flavones (nobiletin, sinensetin and apigenin) as specified earlier in the pre-treatment regime followed by RNA extraction and qRTPCR with primers specific to *cncC* and its downstream target, *gstD1*. We have previously shown induction of both *cncC* and *gstD1* transcripts in response to PQ exposure and used that as the positive control (17). As shown in Fig. 4A, the group of flavones (nobiletin, sinensetin and apigenin) that conferred protection with the pre-feeding mode of treatments failed to upregulate the expression of either *cncC* or *gstD1* in the absence of a stressor like PQ. Next, we determined the effects of flavones (eupatilin, luteolin and tangeretin) in the co-feeding regime along with PQ on the antioxidant genes, *cncC* and *gstD1*. Interestingly, co-treatment with flavones in the presence of PQ resulted in further induction of the *cncC* and *gstD1* transcript levels compared to PQ alone, thereby supporting their role in activating the antioxidant response to counteract against PQ-mediated oxidative stress (Fig.4B). The data suggest that flavones can only activate the antioxidant response pathway in the presence of a stressor like PQ that induces oxidative stress since the pre-treatment feeding regime failed to upregulate the expression of both genes (*cncC* and *gstD1*) involved in the antioxidant response pathway. Based on the differential regulation of antioxidant genes by flavones, our findings suggest that the protective effects of flavones against PQ toxicity are not solely dependent on the antioxidant activities of flavonoids.

**Figure 4.**
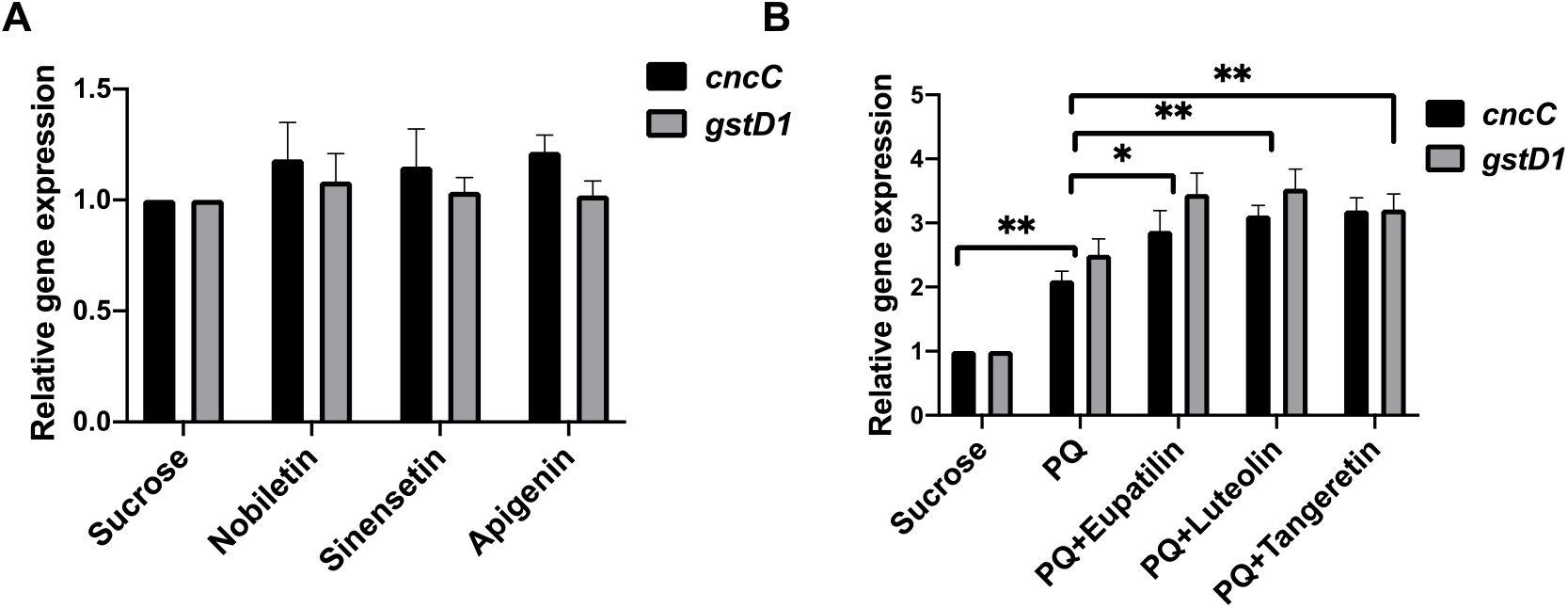
The protective effect of flavones against PQ toxicity is not solely dependent on antioxidant activities. Effects of flavones on the transcript levels of *cncC*, the human Nrf2 orthologue, and its downstream target *gstD1* in *Drosophila* using qRT-PCR. A. Adult male flies were either pre-fed for 4 days with 2.5% sucrose or flavones (nobiletin, sinensetin and apigenin) followed by RNA isolation from the heads of adult flies and processed for qRT-PCR. The transcript levels of *cncC* and *gstD1* were analyzed and plotted after normalization with *rp49* levels as the internal control. Each data point represents mean ± SEM. The mRNA fold changes are normalized to the sucrose-fed (Suc) flies (assigned a value of 1). B. Relative gene expression using qRT-PCR were performed using the co-feeding mode in which adult male flies were exposed to both 5 mM PQ or specified flavones (eupatilin, luteolin and tangeretin) for 24 h. The transcript levels of *cncC* and *gstD1* were analyzed and plotted after normalization with *rp49* levels as the internal control. Each data point represents mean ± SEM. The mRNA fold changes are normalized to the sucrose-fed (Suc) flies (assigned a value of 1). *p<0.05; **p < 0.01 between different feeding conditions based on the Mann-Whitney U test.

### Regulatory effects of flavones on the neuroinflammatory responses associated with PD pathogenesis

Recent studies indicate that chronic neuroinflammatory responses play a critical role in PD pathogenesis (51). Our previous data demonstrate that PQ exposure activates the innate immune transcription factor, Relish, the *Drosophila* orthologue of mammalian NFκB (39). In addition, *relish* knockdown leads to improved survival, climbing ability and rescue of dopaminergic neurons against PQ-mediated toxicity in *Drosophila (39)*. Consistent with these findings, increased NFκB activation has been linked to PD pathogenesis in diverse mammalian models (52-55). Therefore, we next determined the effect of flavones on the transcript levels of *relish*. As observed in earlier studies, *relish* transcript levels were induced 2.64-fold in response to PQ treatment (Fig. 5A). Interestingly, pre-treatment with the following flavones (nobiletin, sinensetin and apigenin) significantly reduced the levels of relish transcripts. The same trend was observed in the co-feeding mode of treatment with the flavones, eupatilin, and tangeretin (Fig.5B). However, luteolin failed to suppress the PQ-mediated induction of *relish* transcripts. It is interesting to note that luteolin was protective at a higher micromolar concentration (100 µM) compared to the rest of the flavones ranging from 1-20 µM. We speculate that it might confer protection by activating the antioxidant response pathways as observed in our study and probably through some other unknown mediators. Our data suggests that flavone-mediated protection against PQ toxicity is dependent on the downregulation of the transcription factor, Relish, thereby supporting the role of flavones as anti-inflammatory mediators.

**Figure 5.**
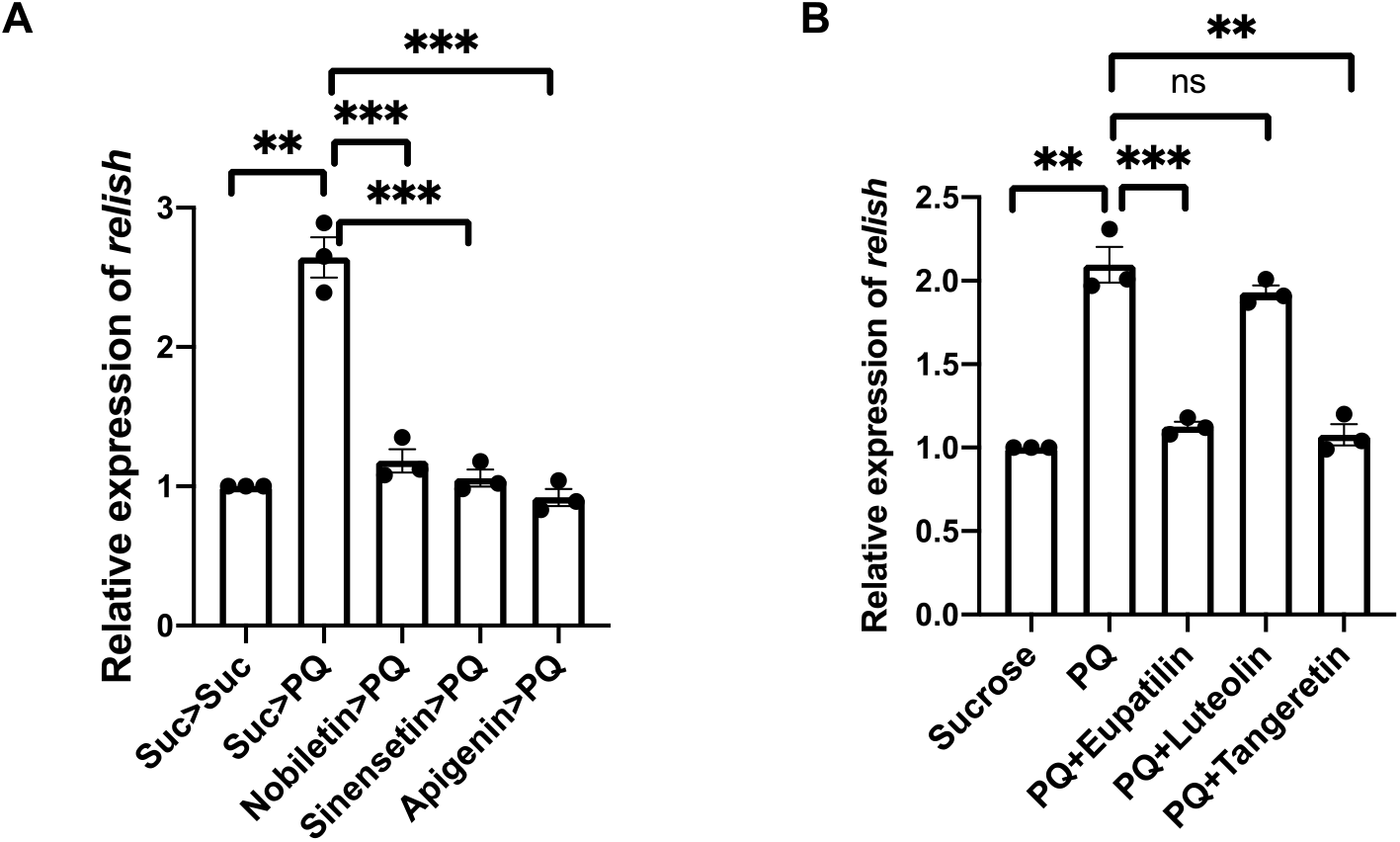
A specific group of flavones confer protection against PQ-induced neurotoxicity through modulation of the neuroinflammatory responses in *Drosophila*. A. Effects of different flavones on the transcript levels of *relish*, the human NFκB orthologue in *Drosophila* using qRT-PCR. A. Adult male flies were either pre-fed for 4 days with 2.5% sucrose or flavones (nobiletin, sinensetin and apigenin) followed by RNA isolation from the heads of adult flies and processed for qRT-PCR. The transcript levels of *relish* were analyzed and plotted after normalization with *rp49* levels as the internal control. Each data point represents mean ± SEM. The mRNA fold changes are normalized to the sucrose-fed (Suc>Suc) flies (assigned a value of 1). \*\**p* < 0.01; \*\*\**p* < 0.001 between different feeding conditions based on Mann-Whitney U test. B. Relative gene expression using qRT-PCR were performed using the co-feeding mode in which adult male flies were exposed to both 5 mM PQ or specified flavones (eupatilin, luteolin and tangeretin). The transcript levels of *relish* were analyzed and plotted after normalization with *rp49* levels as the internal control. Each data point represents mean ± SEM. The mRNA fold changes are normalized to the sucrose-fed (Suc) flies (assigned a value of 1). Statistical differences between the specified feeding groups were analyzed using the nonparametric Mann-Whitney U test \*\**p* < 0.01; \*\*\**p* < 0.001.

## Discussion

In this study, we used toxin-induced *Drosophila* model of PD to investigate the neuroprotective activity of selected dietary flavonoids. Several compounds representing different classes of flavonoids were tested to understand how the structure of these molecules influences their neuroprotective activity *in vivo*. Numerous epidemiological studies have shown that a diet rich in flavonoids decreases the risk of developing neurodegenerative diseases, including PD (4, 8, 11, 14-16, 36). Neuroprotective activity of several flavonoids has also been proven in numerous animal studies, however, the molecular mechanism of action of these compounds remains elusive (2, 12). It also remains unknown whether all classes of flavonoids are characterized by similar neuroprotective activity. Previously, we showed that the polymethoxyflavone gardenin A is a multitarget neuroprotective compound against PQ-induced PD symptoms in *Drosophila* (17). Here a selected group of flavonoids structurally related to gardenin A but differing in the number and positions of hydroxyl and methoxy groups, and the level of saturation of heterocyclic ring C were tested. Our data show that pretreatment with apigenin, nobiletin, and sinsensetin, and co-treatment with eupatilin, luteolin and tangeretin significantly improves survival and mobility defects against PQ-induced parkinsonian symptoms (Figs. 1&3). Interestingly, neuroprotective activities of nobiletin and sinensetin were only observed upon intermittent pretreatment compared to continuous apigenin pretreatment for four days prior to PQ exposure. Previously Kirkland and others have shown that treatment with senolytic drugs, capable of removing senescent cells, was most effective with intermittent administration of these compounds (56, 57). Several polyphenols, including flavonoids have been investigated as potential senolytics, for example: quercetin, fisetin, luteolin, or curcumin (56). The mechanistic nature of this phenomenon is not yet understood and requires further investigation.

All compounds which improved survival and mobility defects in the toxin-induced fly model of PD belonged to one class of flavonoids, namely flavones. Compounds representing other classes of tested flavonoids (flavanones and flavonols as well as all flavonoid glycosides) provided no protection against PQ-induced parkinsonian symptoms. Our data suggest that the presence of α,β-unsaturated carbonyl group, and the lack of substitution at the C3 position are required to confer protection against PQ-mediated toxicity in *Drosophila*.

To the best of our knowledge, this is for the very first time that neuroprotective structure-activity relationships of flavonoids are investigated *in vivo* in the context of a living organism. Previous studies investigating structure-activity relationships for flavonoids have primarily focused on understanding the direct antioxidant activity of these compounds assessed using *in vitro* assays, predominantly in the presence of free radicals (37, 38). Most *in vitro* antioxidant studies indicate that the number and position of hydroxyl groups is the most significant determinant of the antioxidant activity of flavonoids (37, 38). A 3′4′-catechol structure in the B-ring has been indicated as a feature significantly increasing the direct antioxidant activity of flavonoids *in vitro* (37, 38). Additional structural features associated with high free radical scavenging potential *in vitro* include, for example, the presence of a free 3-OH group or the lack of O-methylated groups (37, 38). Many *in vitr*o studies concluded that 2–3 unsaturation and a 4-carbonyl group might be dispensable, although others indicate the presence of these structural features enhances *in vitro* antioxidant activity (37, 38). Ravishankar *et al*. investigated neuroprotective structure-activity relationship in H_2_O_2_-induced oxidative stress *in vitro* model using the SH-SY5Y human neuroblastoma cell line. Structure–activity relationship analysis revealed the following are structural features important for enhancing neuroprotective potential: presence of B-phenyl ring, lack of substitution at C3 position, presence of 4C carbonyl or 4C thiocarbonyl moiety (58). The study also showed that methoxyflavones were less neuroprotective compared to hydroxyl compounds. Our data indicate that both electrophilic α,β-unsaturated carbonyl group and the lack of substitution at the C3 position are essential to confer protection in the PQ-induced model of PD in flies. This data opposes many *in vitro* findings suggesting flavonols, with a hydroxyl group at C3 position are the most biologically active antioxidants (38). Lipophilic flavones with methoxy groups provided better neuroprotection in our model compared to those with hydroxyl groups. This data contradicts the *in vitro* data suggesting the introduction of methoxy groups decreases the biological activity of flavonoids based on in vitro antioxidant assays. However, methylation of flavones was previously shown to enhance their anti-inflammatory activities (59).

Numerous studies, including several clinical trials, disproved the direct antioxidant activity of flavonoids *in vivo* (1). Furthermore, it has been shown that antioxidant activity assessed using *in vitro* assays does not predict antioxidant effects of these compounds *in vivo* (1). *In vitro* studies assume direct interaction between free radicals formed in the cellular environment and flavonoids. This scenario requires high concentrations of antioxidants present in cellular compartments in the close vicinity of free radical production sites. Additionally, many of these studies do not take into consideration the fact that cells contain high concentrations of other molecules involved in antioxidant defense, such as glutathione (1, 2).

The disproval of the direct antioxidant activity hypothesis of natural compounds led to the emergence of other hypotheses explaining the beneficial effects of plant specialized metabolites, including flavonoids. Mattson and his collaborators postulated that many specialized plant metabolites evolved as natural pesticides that protect plants from herbivores and omnivores (1, 18-20, 22). Plants synthesize many compounds that dissuade potential predators from eating plant parts rich in these compounds. Some of these substances are characterized by a bitter taste, such as alkaloids or bitter and astringent tannins (class of polyphenols) (1). To exclude the possibility that any of the tested compounds affects the amount of food ingested by *Drosophila*, we measured the amount of food uptake upon exposure to different flavonoids compared to the control food. Data presented in Fig. 2 clearly indicate that none of the tested compounds significantly changed food uptake. Therefore, the observed protective effects cannot be attributed to other mechanisms like calorie restriction, which has been previously shown to be neuroprotective in multiple animal models (60).

Mattson suggested that most natural compounds are recognized by cells as detrimental and activate multiple evolutionary conserved stress response and other cellular signaling pathways (1). He proposed that natural neuroprotective compounds, including flavonoids act as neurohormetics, providing protection at lower concentrations while exerting detrimental effects at higher concentrations (1, 18-20, 22). Among several molecular pathways, Nrf2 activation has been considered as one of the major mechanisms by which numerous plant specialized metabolites exert beneficial effects (1). We previously showed that antioxidant activity of gardenin A through Nrf2 activation is not sufficient to provide neuroprotection in PQ-induced *Drosophila* model of PD (17). Our data showed that pretreatment with nobiletin, sinensetin and apigenin alone does not result in the upregulation of *Drosophila* Nrf2 orthologue and one of its downstream targets, *gstD1*. This further confirms that antioxidant activity through Nrf2 activation alone is not sufficient to provide protective effects in a toxin-induced model of PD. However, co-treatment with eupatilin, luteolin and tangeretin further increased transcript levels of Nrf2 orthologue and *gstD1*, compared to PQ alone. Our findings suggest flavones increase antioxidant response in the presence of stressors such as PQ. Additionally, our data in the context of the toxin-induced fly model disprove both the neurohormetic and xenohormetic hypotheses, which imply that low concentrations of phytochemicals induce adaptive cellular stress response pathways such as Nrf2 signaling pathway (1, 27).

Following the neurohormesis hypothesis, Mattson suggested that certain phytochemicals should also activate NFκB, similarly to hormetic doses of glutamate and provide protective effects (1). Most known flavonoids inhibit the transcriptional activities of NFkB in cell culture and *in vivo* models and no phytochemicals activating NFκB have been identified to date (61-63). Here we showed that the specific flavones providing protective effects in toxin-induced model of PD, decreased transcript levels of NFκB orthologue in *Drosophila*. Dysregulation of NFκB has been associated with increased neuroinflammation and pathogenesis of several neurodegenerative diseases including PD (64). From among all tested protective flavones, luteolin, failed to decrease transcript levels of NFκB orthologue. Luteolin provided protection at a concentration approximately 5 times higher compared to other flavones. Luteolin is the most polar compound from among all neuroprotective flavones (xlogP values: luteolin – 1.4; nobiletin – 3.0; sinensetin– 3.0; tangeretin – 3.0; eupatilin – 2.9; apigenin – 1.7) with the highest number of free hydroxyl groups. Interestingly, luteolin has been shown to be protective in a transgenic *Drosophila* model of Alzheimer’s disease (65). It is possible that the neuroprotective effects observed for luteolin against PQ-toxicity may be due to the combination of Nrf2 antioxidant effects and its interaction with other unidentified targets.

Lipophilic neuroprotective flavones share similar structural features with anti-inflammatory lipid-derived oxidation products of essential fatty acids (2). Figure 6 compares the structures of anti-inflammatory nobiletin, identified in our study, and the structure of anti-inflammatory 13-EFOX-L_2_ (13-oxoHODE). 13-EFOX-L_2_ is linoleic acid derivative that showed anti-inflammatory effects in a *Drosophila* metainflammation blood tumor model (66). Both molecules represent the class of anti-inflammatory and antioxidant lipophilic soft electrophiles with α,β-unsaturated carbonyl group. Oxidative stress and chronic inflammation have been shown to be underlying hallmarks of numerous noncommunicable diseases, including neurodegenerative ailments (36). Previously, it has been shown that essential fatty oxidation products act as pro-resolving anti-inflammatory molecules through the activation of Nrf2 and NFκB inhibition (67-71). The interplay between the Nrf2 and NFκB pathways has been previously shown to be implicated in the pathogenesis of numerous neurodegenerative diseases (36). It has been proposed that the use of molecules with pleiotropic properties like activating Nrf2 and inhibiting NFκB pathways may emerge as new strategy to combat against inflammatory diseases (72). We postulate that lipophilic flavones along with oxidation products of polyunsaturated fatty acids may constitute a group of pleiotropic soft electrophiles involved in the process of resolution of inflammation through the activation of Nrf2 and NFκB inhibition.

**Figure 6.**
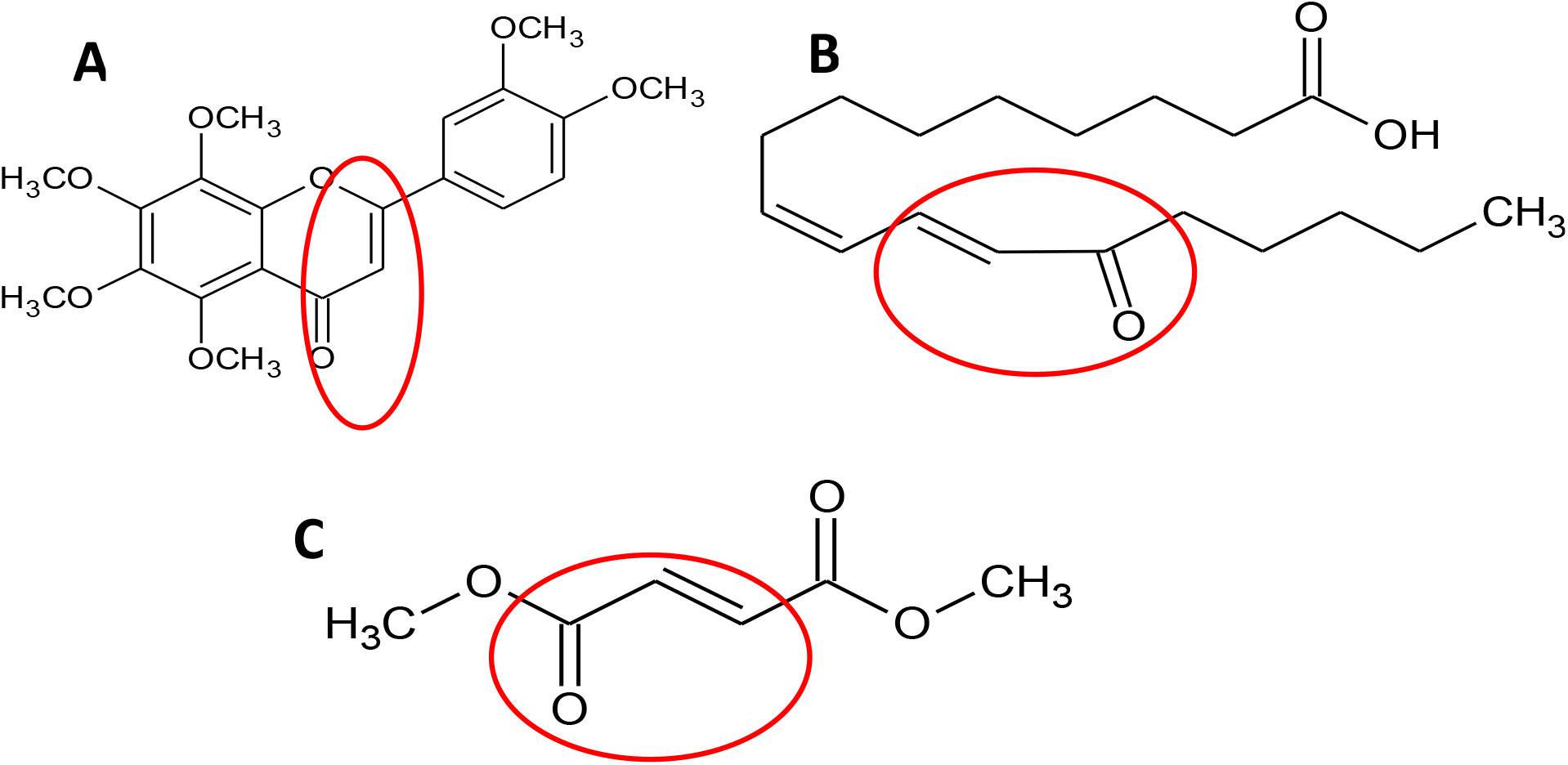
The comparison of structures of soft electrophiles with antioxidant and anti-inflammatory effects. A. Nobiletin (flavone). B. 13-keto-9Z,11E-octadecadienoic acid (13-oxoHODE; 13-EFOX-L2; fatty acid derivative). C. Dimethyl fumarate. Unsaturated α,β-carbonyl group is marked with red.

For a long period of time pluripotent molecules have been disregarded as viable drug leads because of the possible off-target side effects (2, 12). Soft electrophiles, such as omega-3 fatty oxidation products or electrophilic fatty acid nitroalkenes have been also avoided as drug leads due their ability to post-translationally modify numerous protein targets (72, 73). Recent data indicate post-translational modifications of numerous protein targets via electrophilic regulated processes play important roles in the process of resolution of inflammation (72-74). The vast majority of known anti-inflammatory pro-resolving molecules are derivatives of essential fatty acids, previously known as vitamin F (75). Additionally, recent studies have shown that other small molecule soft electrophiles may constitute a viable source of new drug hits and leads for the treatment of numerous diseases (76). For example, dimethyl fumarate, a small-molecule electrophile (Fig. 6C) was recently approved for the treatment of relapsing multiple sclerosis (77). Although molecular mechanism of action of dimethyl fumarate is yet not fully understood, it has been proposed that beneficial effects of this molecule stem from its ability to activate Nrf2 and inhibition of NFκB. Our data show that lipophilic flavones provide protection through activation of Nrf2 and inhibition of NFκB. Similar to other soft electrophilic molecules, phytochemicals with these properties have been disregarded as drug leads (2, 12). The interaction of these molecules with multiple targets led to their classification as improbable metabolic panaceas (IMPs) and pan-assay interference compounds (PAINs) (2, 12). We propose that lipophilic dietary flavones along with oxidation products of PUFAs constitute a class of potentially essential soft electrophiles involved in the process of resolution of inflammation (catabasis). These electrophilic pro-resolving molecules may constitute a class of compounds that should be investigated in the prevention and treatment of multifactorial diseases such as PD. Additionally, we suggest that rather than pursuing these phytochemicals as drugs, we should give them proper consideration as important or even essential nutrients, for example vitamin P, as suggested by Seigler *et al*. that would potentially help to stop or delay the onset and progression of NDs (12).

## Conclusions

To the best of our knowledge, this is the first study that explored neuroprotective structure-activity relationships of flavonoids *in vivo* using the *Drosophila* fly model. Since many *in vitro* studies performed with phytochemicals fail to predict *in vivo* effects, we stress the importance of testing these molecules in a context of a living organism. Our data suggest that the presence of α,β-unsaturated carbonyl group, and the lack of substitution at the C3 position are required to confer protection against PQ-mediated toxicity in *Drosophila*.

Future studies should further explore the concept of essential soft electrophiles and their ability to interact with nucleophiles. Although this study only considered the potential antioxidant and anti-inflammatory properties of flavonoids, other possible targets relevant to PD pathogenesis should be investigated in detail. Similar studies should be extended to additional groups of flavonoids and other classes of phytochemicals in the context of both fly and mammalian models.

## Author contributions

U.M. and L.C. conceived research idea. U.M. designed and performed experiments and analyzed data. J.C. and M.M.O assisted in survival and mobility assays. U.M. and L.C. wrote the manuscript. All authors reviewed and approved the final version of the manuscript.

## Acknowledgements

This work was supported by the start-up funds from the University of Alabama; the National Center for Complimentary and Integrative Medicine of the National Institutes of Health under award number 1R41AT011716-01; NSF-CBET 1915873; NSF-CBET1919906; sponsored research agreement with Wemp LLC. The content is solely the responsibility of the authors and does not necessarily represent the official views of the National Institutes of Health.

